# Neuromesodermal progenitors separate the axial stem zones while producing few single- and dual-fated descendants

**DOI:** 10.1101/622571

**Authors:** Timothy R. Wood, Anders Kyrsting, Johannes Stegmaier, Iwo Kucinski, Clemens F. Kaminski, Ralf Mikut, Octavian Voiculescu

## Abstract

Most embryos and regenerating tissues grow by the action of stem zones. Two epithelial stem zones drive axial elongation in amniotes: the mature organizer generates mesoderm, the neuralised ectoderm around it extends the neuraxis. Bipotential progenitors were also shown to exist. How are these stem cell populations organised and what controls the cell fate of bipotential progenitors? We use direct, in vivo imaging of these stem cells in the chick. We find that progenitors of single and dual fates are mingled in a small region between the specialised stem zones. Divergent tissue movements surround this region. When transplanted downstream of these flows, cells from the region of mixed fates adopt the molecular identity and behaviour of the target stem zone, irrespective of their normal fate. Thus, multipotent cells serve to separate the specialized stem zones, instead of a classical boundary. We propose their fate is determined extrinsically by morphogenetic shearing.

## Introduction

In most animal embryos, only anterior parts are specified during gastrulation; the rest of the body is gradually laid down, in head-to-tail progression, by additions of cells generated by posterior growth zones (Davis and Kirschner, 2000). Vertebrates use this developmental mode at various degrees (Martin and Kimelman, 2009; Wilson et al., 2009): the amniotes to generate the entire main axis except the head (Aires et al., 2018; Henrique et al., 2015), fish and urodeles for posterior trunk and tail extension (Kimelman, 2016; Taniguchi et al., 2017). An historical question, still unresolved, is whether the posterior growth zone is composed of separate mesodermal and neural progenitors (Vogt, 1926), undifferentiated bipotent cells (Holmdahl, 1925), or a combination of these (Holmdahl, 1939).

The establishment of the posterior growth zone is better understood in amniotes, where it marks the end of gastrulation stage. Gastrulation is achieved at the midline primitive streak (PS) by cooperative epithelial-to-mesenchymal transitions (EMT) (Voiculescu et al., 2014). The PS continuously recruits cells from the epiblast lateral to it, replacing the ingressed cells. Cells egressing from the caudal end of the PS form ventral-most mesoderm, with progressively more dorsal types arising from positions further along the PS (Psychoyos and Stern, 1996). During these stages, the anterior PS (Hensen’s node) generates mostly definitive endoderm and acquires organizer properties (Joubin and Stern, 1999).

Two key events establish the posterior growth zones. Firstly, the anterior PS abruptly ceases to recruit cells (Sheng et al., 2003) and starts regressing caudally. The region acts as a mesodermal stem zone (MSZ), harbouring long-term resident progenitors that generate the internal dorsal tissues (Selleck and Stern, 1992a): prechordal mesendoderm and notochord from its front sector, and medial paraxial mesoderm from the sides (Iimura et al., 2007; Selleck and Stern, 1991, 1992b) (see Figures 1a and 1b; Supplementary Video S1). Secondly, the epiblast around the MSZ acquires definitive neural character. Most of the neural territory, lying anterior to the node, contributes to brain (Figures 1c and 1d; Supplementary Video S2). The regions lateral to the anterior PS become neural stem zones (NSZ), which then elongate the neuraxis caudal to anterior hindbrain (Brown and Storey, 2000; Mathis et al., 2001). Similar fate maps were drawn in the mouse embryo (Kinder et al., 2001).

**Figure 1.**
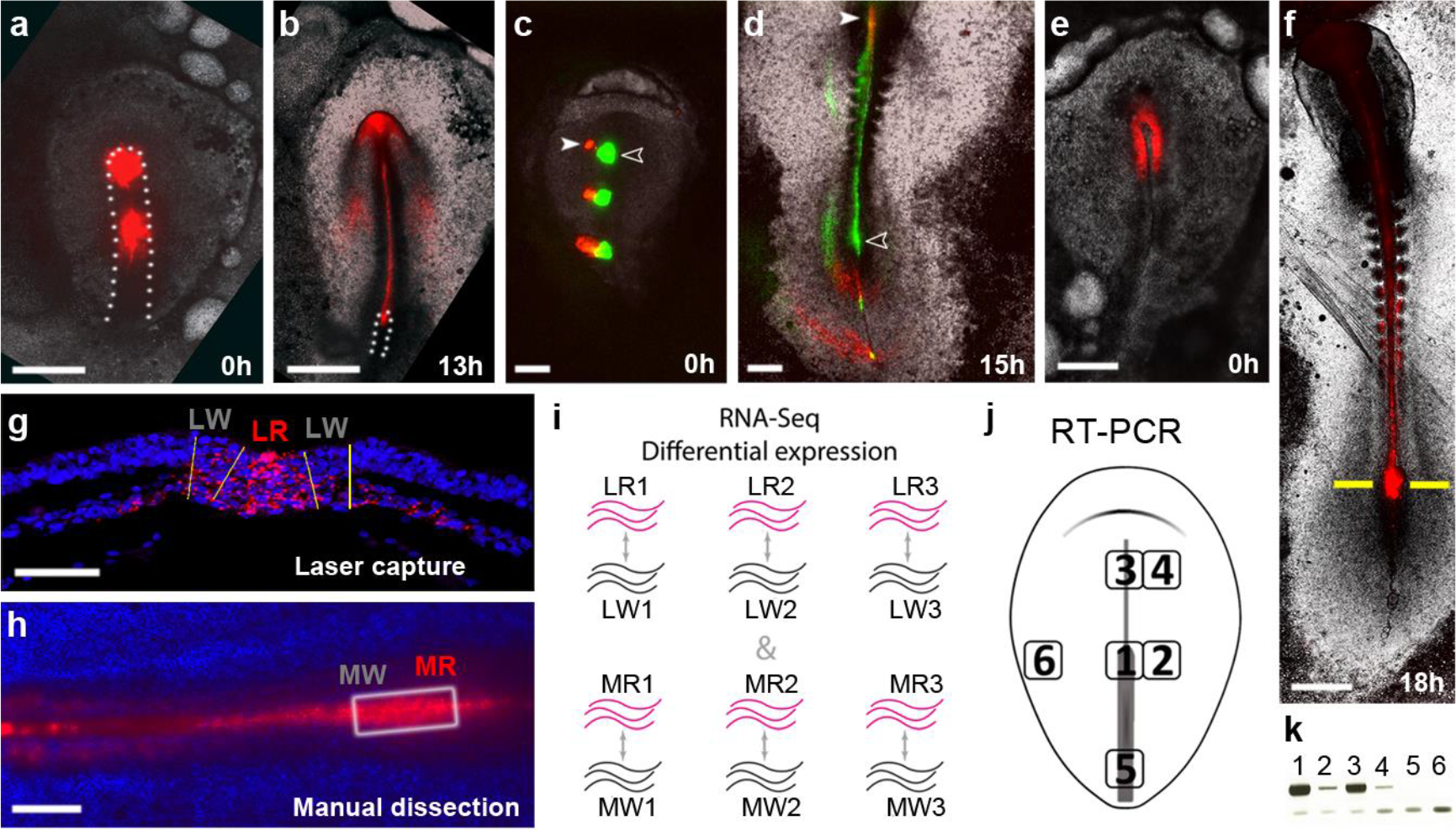
Two distinct stem zones drive axial elongation, allowing separation of their cells. **a, b** The tip of the primitive streak (PS, dotted outline) becomes a stem zone for dorsal mesoderm. The anterior and middle of the primitive streak (PS) labelled fluorescently (red) at the end of gastrulation (**a**). Only the anterior PS (MSZ) retains long-term resident cells, as a reservoir of axial and paraxial mesoderm (**b**). **c, d** The epiblast around the anterior MSZ becomes neural instead of contributing to mesoderm. The anterior, middle and posterior PS (green fluorescence), and adjacent epiblast positions (red fluorescence) labelled at the end of gastrulation (**c**). Cells lateral to the node (solid arrowhead) contribute to neural plate, while the MSZ generates axial and paraxial mesoderm (open arrowhead, **d**). In contrast, more caudal epiblast ingress through the mid- and caudal PS and form mesoderm. **e, f** Exemplar embryo used for differential gene expression between the MSZ and surrounding regions. **e** Fluorescent labelling of the anterior PS at the end of gastrulation. **f** Embryo in after culture, used to isolate resident cells in the MSZ. Line indicates the level of sections used in laser-capture micro dissection (LCM) in (**g**). **g** LCM of resident cells in the MSZ (red fluorescence, LR) and their unlabelled neighbours (LW). **h** Caudal part of an embryo similar to (F); body axis is horizontal, anterior to the left. Boxed, region excised to manually dissect MSZ cells (MR) and their neighbours (MW). **i** Scheme for differential expression of RNA-seq profiles between the MSZ and neighbouring cells. **j, k** Genes with higher gene counts in the MSZ checked for specific expression by RT-PCR. **j** Regions used to generate cDNA. **k** Results for a gene (*cTPO*) typically selected for *in situ* hybridization screen. Scale bars, 500 μm (a-f); 200 μm (g, h).

It has long been debated whether these specialised stem zones are clonally separated or common, neuro-mesodermal progenitors (NMPs) exist (Catala et al., 1995; Le Douarin et al., 1998; Patten et al., 2003). In chick, labelling of single (Brown and Storey, 2000; Selleck and Stern, 1991) or small groups of cells (Brown and Storey, 2000) found very few cells of double progeny in the vicinity of the anterior PS. Such progenitors were also deduced to exist in mouse from retrospective clonal analyses (Tzouanacou et al., 2009). However, neither this approach nor extensive lineage reconstruction from live imaging datasets (McDole et al., 2018) could map the NMPs.

Molecularly, the neural domain is marked by the expression of *Sox2* (Sheng et al., 2003), which marks only neurally-fated cells (Mugele et al., 2018) and is sufficient to neuralise even nascent mesoderm if ectopically expressed (Takemoto et al., 2011). A molecular identity of the MSZ, also committed to its fate, is lacking. However, it is and expresses generic pro-mesodermal genes (*Bra*, *Tbx6*), which are in a mutually repressive relationship with *Sox2* (Gouti et al., 2017; Javali et al., 2017; Koch et al., 2017; Takemoto et al., 2011). A few cells co-expressing pro-neural and pro-mesodermal genes were identified around the organizer in chick (Henrique et al., 2015), mouse (Garriock et al., 2015; Tsakiridis et al., 2014), fish (Martin and Kimelman, 2012) and frog (Davis and Kirschner, 2000; Gont et al., 1993). Also, some *Bra*^+^ cells in mouse have neural destiny (Garriock et al., 2015). Transplantation experiments showed that regions from around the mouse node can contribute to both neural and mesodermal tissues (Cambray and Wilson, 2002, 2007; Wymeersch et al., 2016). Pluripotent stem cells from both mouse and human can be induced to co-express this antagonistic set of genes, and produce either mesoderm, neural or neural crest cells (Denham et al., 2015; Frith et al., 2018; Gouti et al., 2014; Lippmann et al., 2015; Tsakiridis et al., 2014; Turner et al., 2014). These suggest that, at population level, cells of neuromesodermal potential correspond to, or are a subset of the *Sox2*^+^, *Bra*^+^ population, although this definition has recently been challenged (Mugele et al., 2018). The normal fate of bipotent cells could not be established.

Contrasting views of the posterior stem cell populations exist. On the one hand, mechanisms were described that could ensure the segregation of NSZ and MSZ cells (Acloque et al., 2011, 2012, 2017; Sheng et al., 2003). On the other hand, it was suggested that the bipotential cells control production of neural and mesodermal tissues, via the balance of *Sox2* and *Tbx6*/*Bra*. This could be altered either in response to extrinsic signals (Martin and Kimelman, 2012) or by the natural evolution of the gene regulatory network (Gouti et al., 2017).

Key questions remain: What are the location of bipotential progenitors and their spatial organisation? How do they divide? What are their respective contributions to axial elongation? What role do they play in development? Here, we use molecular profiling to find ways to distinguish clearly between the specialised stem zones. We then present a comprehensive cell lineage analysis of the stem zones involved in axial elongation, based on direct observations of the stem zones in live chicken embryos, and experimentally explore the mechanisms of lineage segregation.

## Results

### A unique molecular profile of the mesodermal stem zone

While the transcriptomes of the neural stem zones and of the putative neuromesodermal precursors have been explored (Gouti et al., 2017; Verrier et al., 2018), the mesodermal stem zone is poorly characterised. As a prerequisite to assess the molecular identity of cells in posterior stem zones in functional experiments, we first sought to identify a specific molecular identity of MSZ, distinct from both the surrounding neural plate and the more caudal PS that continues to gastrulate. We fluorescently labelled (CM-DiI) the organizer epiblast just prior to it becoming a stem zone (at fully extended PS, stage HH 3^+^; see Figure 1e); embryos were then left to elongate to 8-10 somites (Figure 1f). Cells remaining in the MSZ were collected and, separately, their neighbours in the neural plate and emerging paraxial mesoderm. Two methods were employed: laser capture microdissection from fixed, sectioned specimens and manual separation from live embryos (Figures 1g and 1h). For each method, three pairs of biological replicates were deep sequenced and compared (Figure 1i and Materials and Methods). We selected 93 transcripts showing consistent and higher expression in the MSZ (see Supplementary Table S3). For an initial assessment of their specificity of expression, we used RT-PCR on cDNA from pieces of tissue dissected from the MSZ and five other embryonic locations (Figure 1j and 1k). Thirty-two genes (Supplementary Table S4) showed restricted expression in the MSZ and the midline structures anterior to it; their expression was further examined by in situ hybridization. This revealed 4 genes that are turned on in the anterior PS at the end of stage HH 3^+^, and afterward remain expressed there and in the tail bud: *cWIF1*, *cTPO*, *cPTGDS* and *cUCKL1* (Figures 2a-p). Their expression is similar in the anterior PS, and is absent from the presomitic mesoderm originating laterally from the anterior PS; expression persists in the emerging notochord, to different degrees between these genes. Transcripts were not detected in any region of the neural plate.

**Figure 2.**
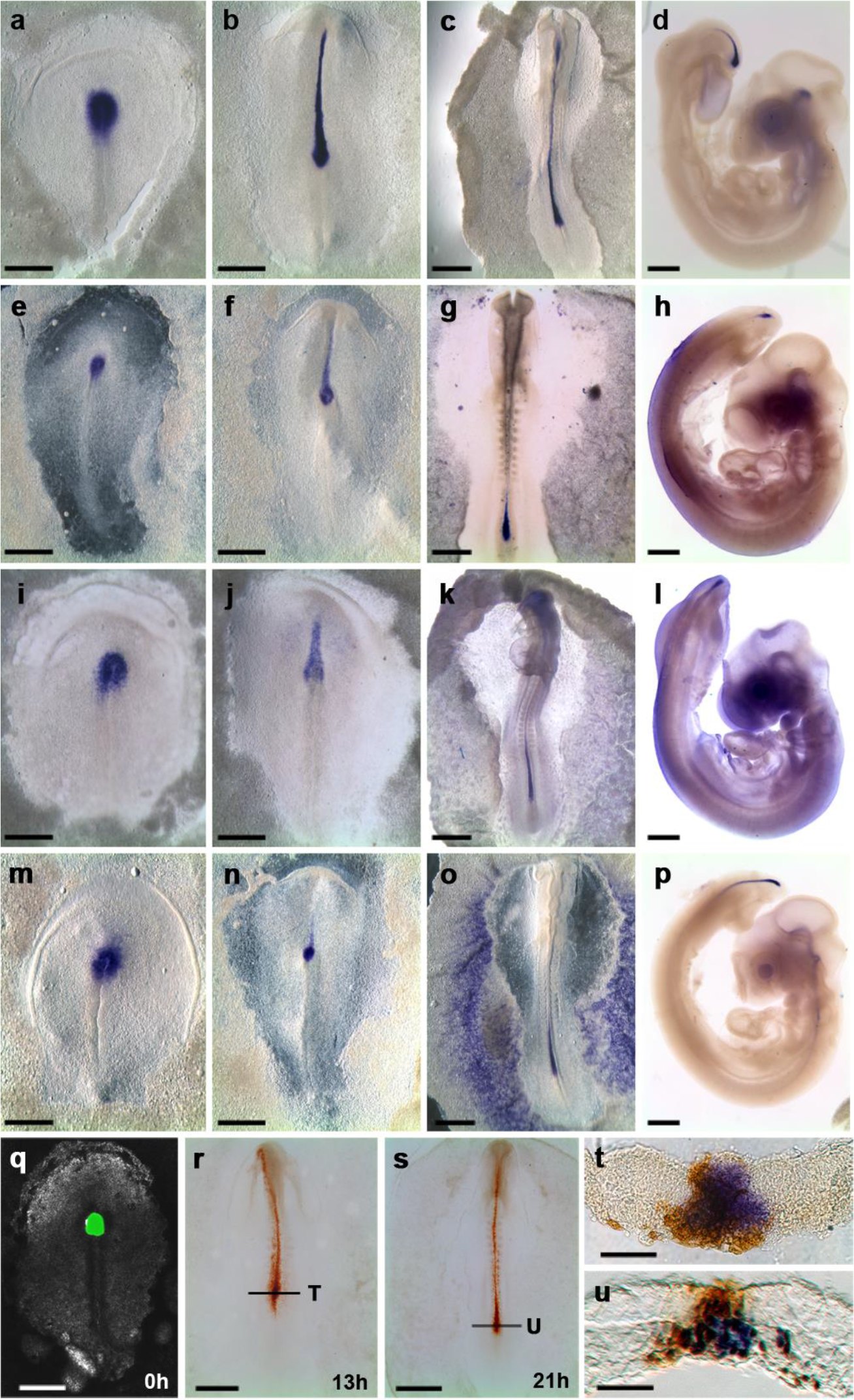
The mesodermal stem zone displays a constant molecular signature. **a-p** Four genes upregulated only in the mesodermal stem zone at its formation, and remain expressed at subsequent stages: *cWif1* (**a-d**), *cThPO* (**e-h**), *cUCLK1* (**i-l**), *cPTGDS* (**m-p**). **q** Whole organizer (green fluorescence) grafted homotopically into an unlabelled embryo at the end of gastrulation. **r, s** At subsequent stages, all descendants of the labelled cells (brown) residing in the NSZ express *cWif1* (purple). **t, u** sections at the levels indicated in (R, S), respectively. Scale bars, 500 μm (a-c, e-g, i-k, m-o, q-s); 1 mm (d, h, l, p); 100 μm (t, u).

Do these transcripts faithfully mark the MSZ at all stages? To label all cells of the MSZ, we grafted anterior PS regions homotopically and isochronically, at the onset of elongation (late stage HH 3^+^), from embryos labelled fluorescently (CMFDA) into unlabelled hosts (Figure 2q, n=25). At subsequent stages (5-15 somites), we found that long-term resident cells (remaining in the MSZ) express all four genes (Figures 2r-u).

Wif1 is an antagonist of both canonical and non-canonical Wnt ligands (Hsieh et al., 1999). The prostaglandin synthase PTGDS was implicated in controlling EMT (Zhang et al., 2006) and transporting retinoids (Tanaka et al., 1997). Thrombopoietin controls proliferation in the hematopoietic niches (de Graaf and Metcalf, 2011; Hitchcock and Kaushansky, 2014; Kaushansky, 2016), and UCKL1 in blast transformation (Kashuba et al., 2002; Michaille et al., 2005). Importantly, none of these proteins is known to regulate any of the other three; indeed, their genes do not belong to known synexpression groups, such as those shared by different embryonic organizers (Anderson et al., 2016). Therefore, their co-expression provides a unique MSZ molecular identity.

### *In vivo* imaging of the axial stem zones reveals the dynamics of axial stem cell lineages

In parallel, we asked where the progenitors of the specialised stem zones arise, whether they are separated or mixed, if and where common progenitors exist. To capture the dynamics of the process, the epiblast regions fuelling axial elongation and their surroundings were directly imaged, using high-resolution multi-photon time-lapse microscopy (Voiculescu and Stern, 2012) in live transgenic chick embryos expressing membrane-tethered EGFP ubiquitously (Rozbicki et al., 2015) (Figures 3a and 3b). The fifteen-hour sequence encompasses the final stages of gastrulation (HH 3^+^) and axial elongation up to the formation of 10 somites (HH 10), similar to the embryo in Supplementary Video S2 (Supplementary Videos S5 and S6). This approach permits both prospective and retrospective lineage analyses. Automatic segmentation and cell tracking (Stegmaier et al., 2016) correctly identified more than 95% of cells between cell divisions; this approach provides a view of the cellular flows at high density. To gain perfect accuracy in establishing lineage relationships, we manually tracked 752 lineage trees, with 732 cell divisions; this manually annotated dataset includes virtually all cells at the interface between the neural and mesodermal stem zones. Agreement between the cell movements obtained with these approaches is shown in Supplementary Video S7. The cell movements and behaviours characteristic for each epiblast domain are summarised below.

**Figure 3.**
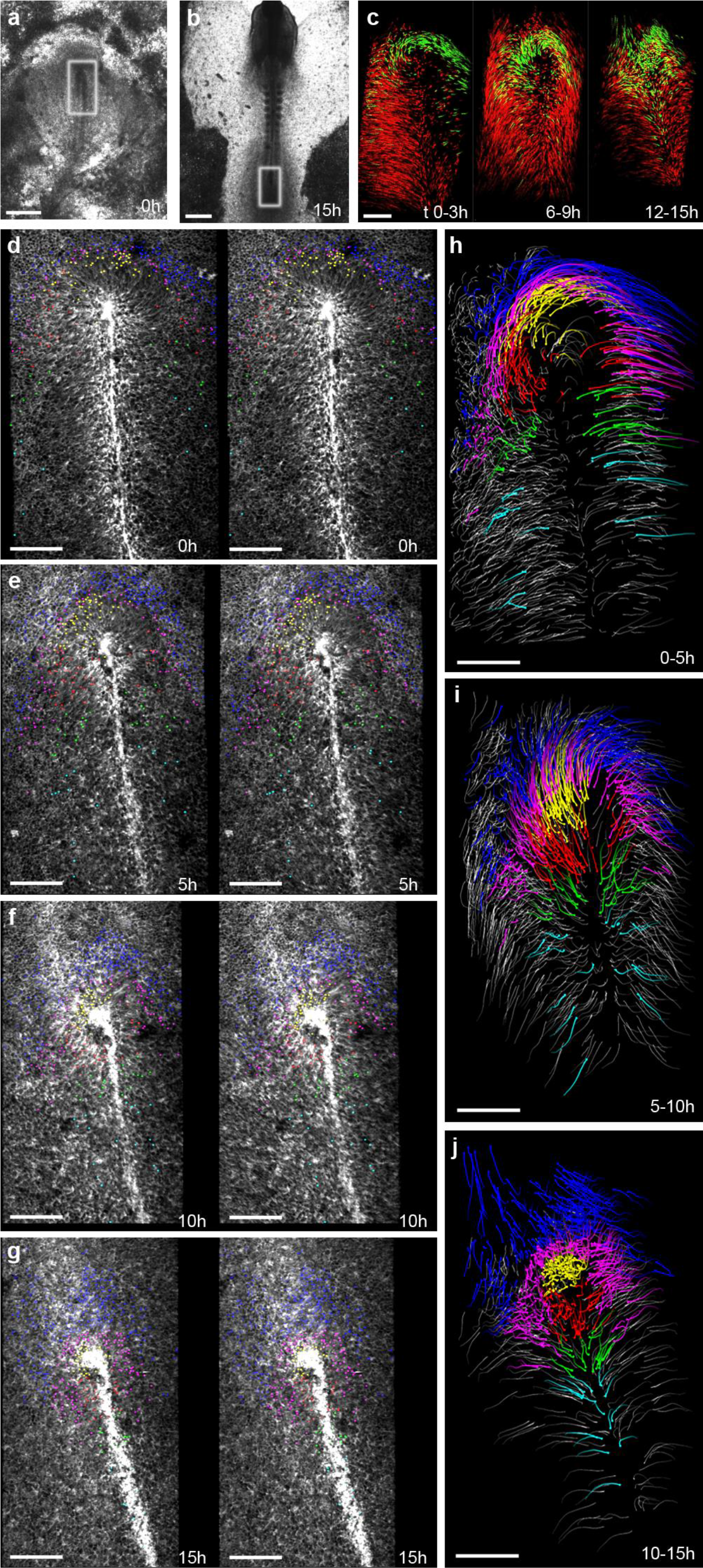
Direct *in vivo* imaging of the stem zones allows they dynamic mapping of progenitors fuelling axial elongation. **a, b** Multi-photon imaging of the stem zones in a membrane-GFP transgenic chick embryo. Embryo at the end of gastrulation (**a**) and after 15 hours of imaging (**b**); the region imaged at higher resolution is boxed. **c** Comparison between trajectories of automatically and manually tracked cells (red and green, respectively). Imaged volumes are registered with respect to the centre of the organizer. **d-f** Location of manually tracked cells contributing to the neural plate (blue), the mature organizer (yellow and red), and regions of the primitive streak posterior to it (green and turquoise); stereo pairs at the times indicated. Cells in magenta do not move into either the MSZ or the neural plate during the sequence. **h-j** Trajectories of cells displayed in (**d-f**) during the indicated intervals. A few automatically tracked cells are shown in grey. Scale bars, 500 μm (a, b); 100 μm (c-i).

In the PS caudal to the node, cells ingress by epithelial-to-mesenchymal transition (EMT) within 2 hours of entering the PS. They are replaced by cells from the lateral epiblast that move in bilaterally symmetrical trajectories towards the PS, similar to gastrulation stages (Voiculescu et al., 2014) (green and turquoise in Figures 3d-g and 3H-J, Supplementary Videos S8 and S9). Most cells present in the anterior PS at the end of gastrulation also ingress fast. They are replaced by the long-term resident cells originating from the periphery of the node at the end of gastrulation (yellow and red in Figures 3d-g and 3h-j, Supplementary Videos S8 and S9), and composing the Hensen’s node at the end of the sequence. During the first 5 hours after the end of gastrulation (HH 4-6), the cells in the right-hand sector of the node transiently rotate by about 30° leftward as they move toward the centre of the PS (Figures 3h-j), confirming earlier observations (Gros et al., 2009; Mendes et al., 2014). The resident cells occupying the very tip of the PS at late stages (yellow) are derived from cells located in an anterior ∇-shaped quadrant of the organizer, previously shown as the origin of prechordal mesendoderm and notochord (Selleck and Stern, 1991). The remainder of the resident cells (red), which give rise to medial somitic mesoderm (Iimura et al., 2007; Selleck and Stern, 1991), originate from the lateral sides of the Hensen’s node at the end of gastrulation.

Neural plate-destined cells (blue) were annotated based on their position. Instead of moving towards the PS (as the cells in the epiblast lateral to it) or the NSZ, they persistently translocate anterior to the node, and none is then seen moving in the opposite direction. Some cells could not be classified as belonging to either the NSZ or the MSZ domain from their movements during the sequence or from their final location (magenta in Figure 3 d-j); we refer to their domain as the “zone of neuromesodermal precursors” (ZNMP).

We found two types of lineages where sister cells have different fates. These are presented separately below.

### Individual cells undergo EMT and ingress from the neural plate

Cells arriving at the PS caudal to the NSZ ingress with high probability (Figure 4a), none dwelling for longer than 2 hours (Supplementary Video S10). However, these cells undergo EMT singly, being surrounded by cells forming rosettes (see Figure 4d), and which ingress later. Surprisingly, we observed infrequent EMT events outside the PS: in the epiblast lateral to the PS, the ZNMP between the neural and the mesodermal stem zones, and all regions of the neural plate (Figure 4b and 4c, Supplementary Video S10). These EMT events are qualitatively indistinguishable from those in the PS (Figures 4e-g), but occur with very low probability (1.6% per cell per hour). Of note, the sisters of ingressing cells do not ingress at the same time, and most persist throughout the sequence in their neural or ectodermal territories.

**Figure 4.**
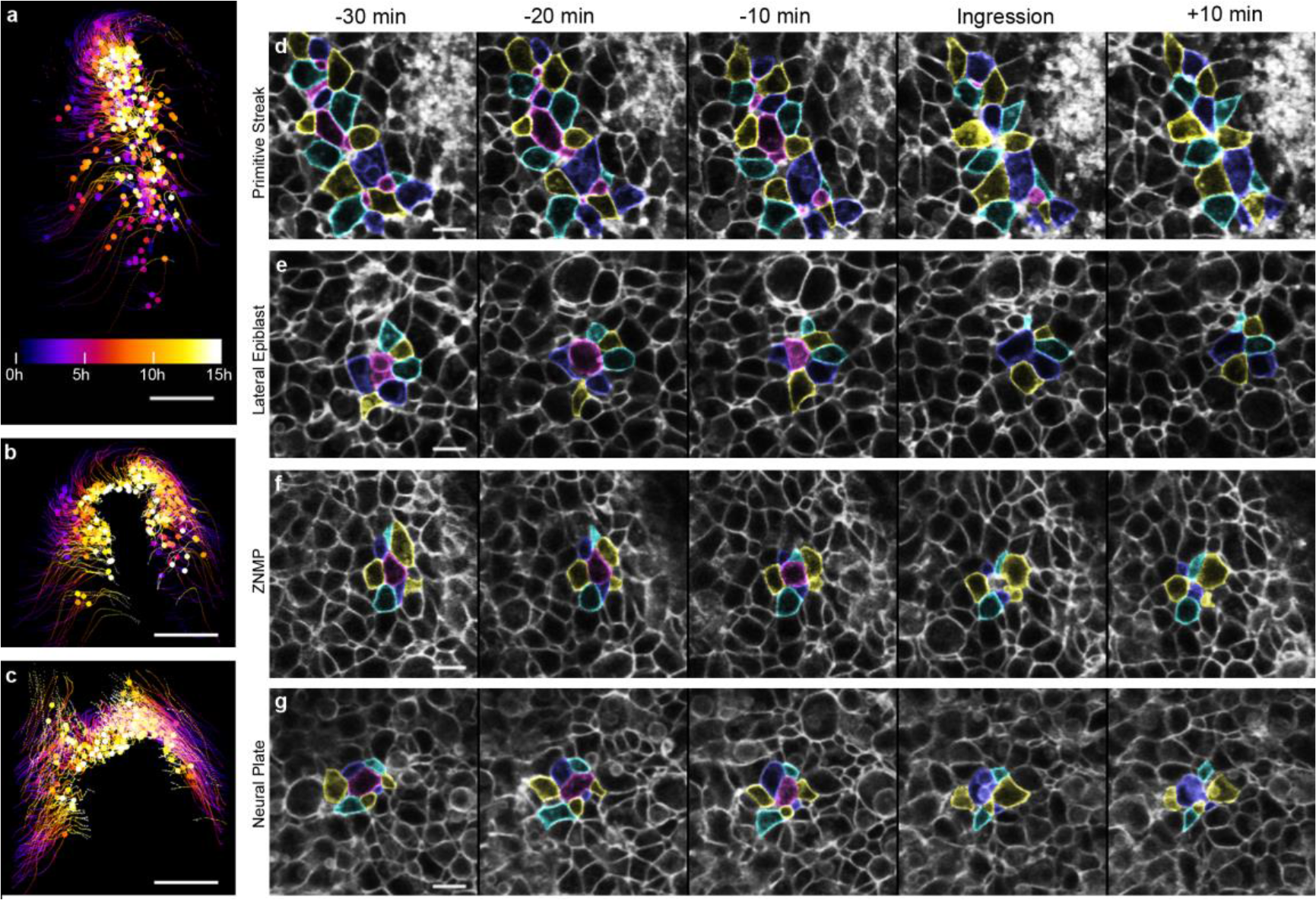
Individual epithelial to mesenchymal transitions occur in the neural and non-neural epiblast during elongation stages. **a-c** Location of EMT (ingression) events during axial elongation in the PS and epiblast lateral to it (**a**), the zone of neural and mesodermal precursors (**b**) and the neural plate (**c**). The time of occurrence is colour-coded. **d-g** Sequence of EMT events. Ingressing cells are coloured magenta and surrounding cells yellow, cyan and blue. Examples are shown in the primitive streak (**d**, midline on the along the right edge of panels), epiblast lateral to the PS (**e**), the zone of neural and mesodermal precursors (**f**) and the neural plate (**g**). Scale bars, 100 μm (a-c); 10 μm (d-g).

### A narrow domain contains long-term bipotential cells mingled with precursors of single fate

When tracking neural and node cells back to the end of gastrulation, we observed a crescent-shaped region of around 200 cells where precursors of the late organizer (MSZ) are mingled with neural plate progenitors, including the NSZ (magenta in Figure 5 a-c, and Supplementary Videos S8 and S11). During the 15 hours of axial elongation examined here, most of the cells in this region (ZNMP) do not move into either the late organizer or the neural plate. Only 15 progenitor cells produce daughters of different fate (Figure 5 a-c and Supplementary Video S11): one produces descendants in both the neural and mesodermal stem zones, and the rest have one daughter in the NMPZ and another one in a committed domain (6 in MSZ and 8 in the NSZ).

**Figure 5.**
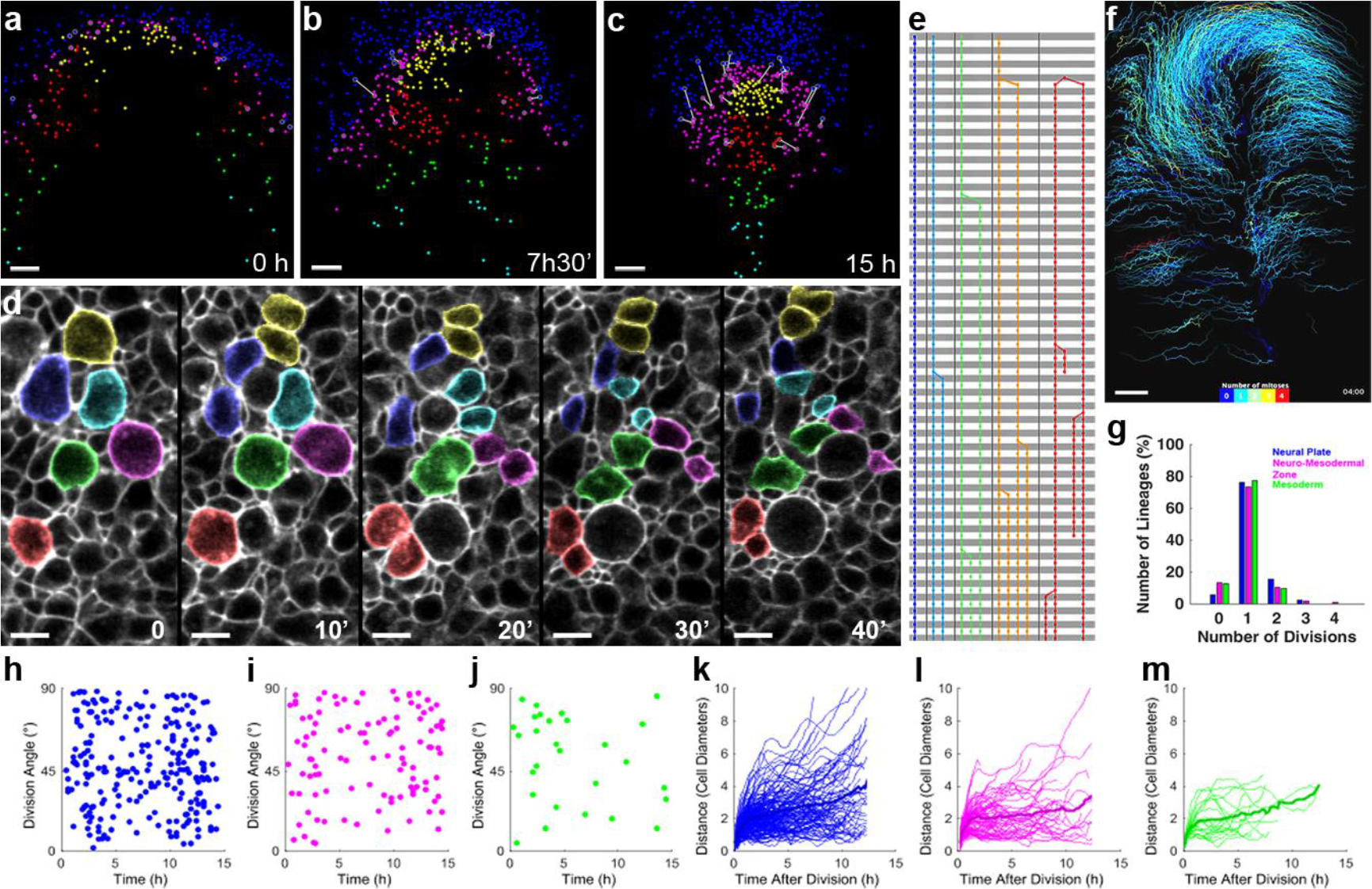
The cells in the neuromesodermal region display the same modes of cell division and daughter dispersal as those in the neighbouring, specialised stem zones. **a-c** A subset of cells in the ZNMP (magenta) have progeny of different fates. Sister cells are connected by white bars. Cells coloured according to fate: the neural plate – blue; node sectors generating notochord and somatic mesoderm - yellow and red, respectively; more caudal PS - green and turquoise. **a** Neuromesodermal precursors cells are initially mingled with precursors of only neural and mesodermal fates. **b, c** Lineages separate over time. **d** Sister cells (coloured) disperse by different amounts following cell division: some remain in contact (yellow, red), but most separate immediately (blue, turquoise, green, magenta). **e** Examples of lineage trees, coloured by the number of cell divisions observed. Time runs from top to bottom, in 10’ ticks. **f** Lineages with different number of mitotic events, color-coded as in **e**, are uniformly distributed. **g** The distribution of lineages with different number of cell division is similar in the neural, neuromesodermal and prospective mesoderm territories. **h-j** Orientation of cell divisions with respect to direction of mother cell in the neural plate (**h**), neuromesodermal zone (**i**) and mature organizer (**j**) lineages. **k-m** Distance between daughter cells in the neural plate (**k**), neuromesodermal zone (**l**) and mature organizer (**m**) domains; the mean is shown as a thick line. Scale bars, 50 μm (a-c, f); 10 μm (d).

### All cells in the ZNMP divide isotropically and at the same rate as in the neighbouring regions

Do cells of different fates (fated to become neural, mesodermal or in the ZNMP) have different modes of cell division? In all three lineages, dividing cells follow the same sequence of morphological changes (Figure 5d): the mitotic cell rounds up at the apical surface, compressing the neighbours; the daughter cells usually separate, although some remain in contact (see below), and cause rearrangement of neighbouring cells, confirming and extending previous observations (Firmino et al., 2016). The number of divisions observed within 15 hours varies between 0 and 4; asymmetry is seen with respect to the length of the cell cycle within each family tree with multiple divisions (Figure 5e). These types of lineages are uniformly distributed spatially, irrespective of cell fate (Figure 5f and Supplementary Video S12), and have the same frequency in the populations destined to become neural, mesodermal, or both (Figure 5g). No preferred orientation of cell division planes was detected with respect to direction of movement of their progenitors (Figures 5h-j and Supplementary Video S13) or the main axis of the embryo (not shown). Daughter cells separate after mitosis to various degrees and further drift apart, with no significant difference between regions (Figures 5k-m and Supplementary Video S13).

### All cells in the ZNMP are multipotent, irrespective of their normal fate

Since cells in the ZNMP do not differ in their rate or mode of cell division from those of neighbouring regions, what drives cells in the ZNMP to move and differentiate differently from one another? One hypothesis is that the cells of the ZNMP are intrinsically committed to different fates and move accordingly. Alternatively, it could be that all cells in the ZNMP are multipotent and only adopt their fate once they arrive in the neighbouring, committed domains. We observed that the ZNMP sits between divergent morphogenetic movements in the MSZ and the neural plate (Figures 6a and 6b, Supplementary Movie 14), and the occasional cells leaving the ZNMP follow these collective flows. To distinguish between the hypotheses above, we transplanted groups of cells from the ZNMP (Figure 6c) downstream of the collective cell flows surrounding their original location, in the locations where they could end up in 3 hours’ time (see Table1 for a summary of operations). The epiblast corresponding to the ZNMP was transplanted after careful removal of underlying mesoderm; completely to exclude from analysis any residual mesodermal cells, the epiblast of the donor embryo was electroporated with a control, fluorescently-labelled morpholino, and we analysed the molecular identity and contribution of these labelled cells.

**Figure 6.**
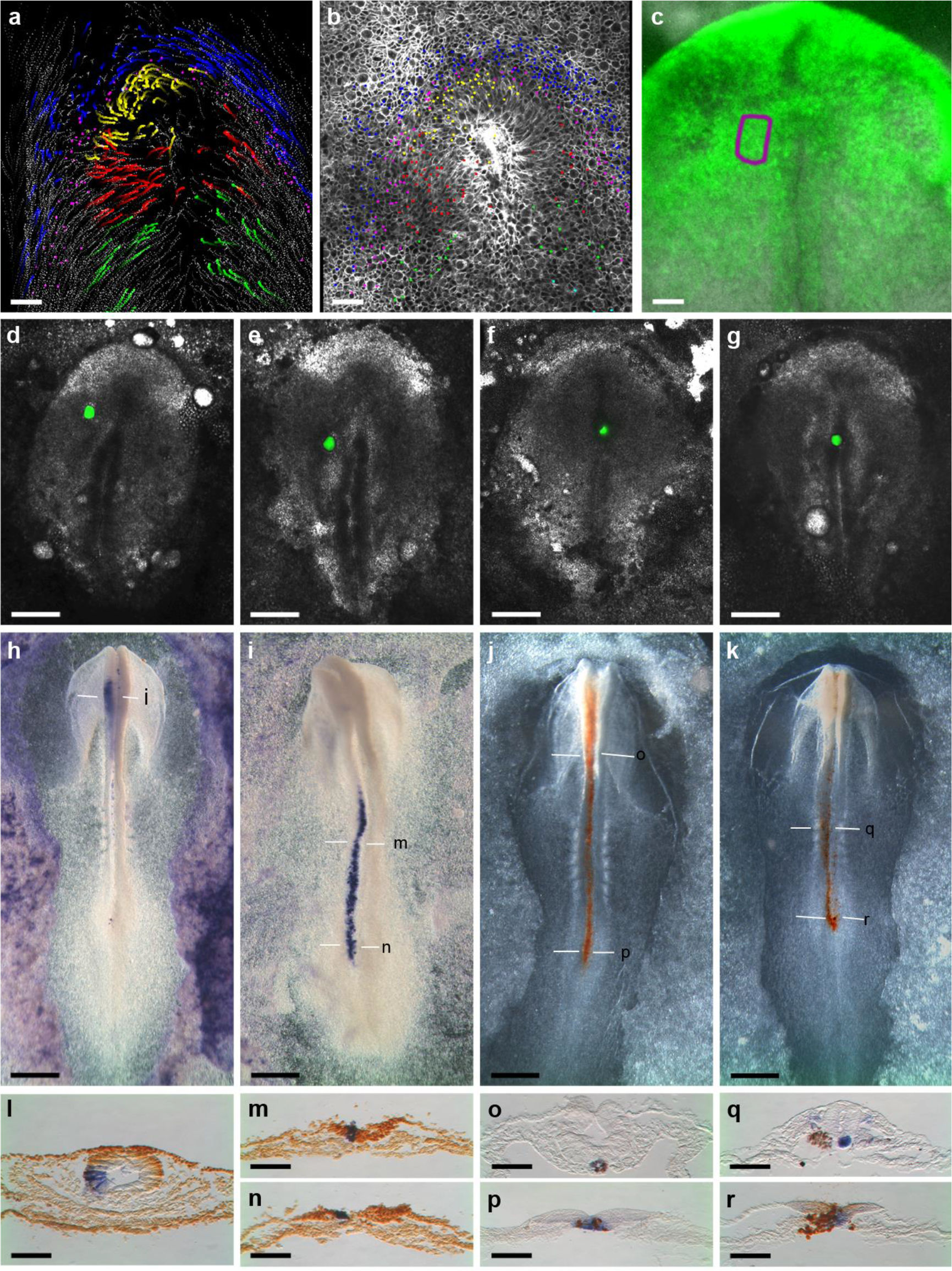
All cells in the zone of mixed precursors are competent to become incorporated into either specialised stem zone. **a** Location of cells in the zone of neuromesodermal precursors (magenta) at stage HH 5. Surrounding cells are coloured blue (neural plate), yellow and red (the mature organizer), green and turquoise (later ingressing through of the more posterior primitive streak). **b** Cells in the zone of neuromesodermal precursors shown against the displacements of neighbouring cells over the following three hours. Trajectories intensify with time, and eventual positions are marked by a dot. A subset of automatically tracked cells (grey) are also shown. **c** Boxed, transplanted region of mixed precursors from a donor embryo of same stage (HH 5). Ectoderm cells are electroporated with a fluorescently-tagged control morpholino. **d, g** Embryos having received quarters of the region boxed in **c** into the neural plate above the level of the node (**d**), neural stem zone (**e**), anterior or lateral mature organizer (**f, g**, respectively). **h-k** The operated embryos in (**d-g**), respectively, after 15 hours of development. Grafted cells are stained purple (**h, i**) or brown (**j, k**). **l-r**, Transversal sections of the embryos above in (**h-k**), at the levels indicated by lines. **l-n** Sox2 immunostaining (brown). **o-r** *cWif1* in situ hybridization (purple). Scale bars, 50 μm (**a-c**, **l-r**); 500 μm (**d-k**).

**Table 1.** Summary of ZNMP grafts. Total successful operations, in which the grafts integrated into the target tissue. Numbers in brackets indicate the total operations performed.

|  |  | Stage of Operation |  |  |
| --- | --- | --- | --- | --- |
|  |  | HH 4 | HH 5 | HH 6 |
| **ZNMP grafted into** | **Definitive neural plate** | 12 (25) | 13 (30) | 10 (24) |
|  | **Neural stem zone** | 14 (25) | 15 (30) | 9 (24) |
|  | **Anterior node** | 18 (25) | 20 (30) | 15 (24) |
|  | **Lateral node** | 20 (25) | 19 (30) | 17 (24) |

When grafted into the neural domain anterior to the node (Figure 6d), all cells contributed to a small region of the brain (Figures 6h and 6l). When brought into the neural stem zone lateral to the node, all cells contributed to a long stripe along the more posterior neural plate (Figures 6i, 6m and 6n), with some residing in the NSZ. Conversely, all ZNMP cells transplanted into the axial- or paraxial mesoderm-producing regions of the MSZ (Figures 6f and 6g, respectively) gave rise to the normal derivatives of those regions, some becoming residents of the MSZ (Figures 6j and 6k). In such experiments, it is possible that grafted cells were simply carried away by adhering to the neighbouring host cells. We tested this alternative and found that the transplanted ZMP cells remaining in the mesodermal stem zone also upregulated *cWIF1* (Figures 6o and 6p for the notochord region, and Figures 6q and 6r for the somitogenic region of the MSZ) and the other MSZ genes (not shown), as appropriate for this stem zone.

## Discussion

Previously, two committed stem zones (mesodermal, MSZ, and neural, NSZ) were shown to drive axial elongation after gastrulation; however, indirect evidence also indicated that they share some common, neuromesodermal progenitors (NMPs) during elongation (see Introduction). These could not be directly observed and mapped in mouse (McDole et al., 2018), chick or zebrafish (Attardi et al., 2018) at elongation stages. There is also some controversy regarding their molecular definition and the interpretation of genetic manipulations at population level to deduce the individual cell behaviours fate decisions that allocate cells to the specialised stem zones (Mugele et al., 2018). Here, we used direct, in vivo imaging and comprehensive lineage tracing to determine the relationships between the axial stem zones in the chick.

Our data show that individual cells produce progeny of dual fate by two distinct processes, occurring in different embryonic domains. One source are rare EMTs (frequency of ~1,6% per cell per hour) occurring outside the primitive streak, including the neural stem zone and the differentiated neural plate (Figure 4 and Supplementary Video S8). Every cell undergoing EMT has a sister either ingressing at a different time-point or, in the majority of cases, remaining in the ectoderm during the duration of the sequence. The ingressed cells may insert into the underlying mesoderm (pre-somitic mesoderm or notochord), where they could give rise to clones of short antero-posterior extent. Their siblings, remaining either in the neural stem zone or the differentiated neural plate, are likely to contribute long and, respectively, short clones in the neural tissue. We propose this explanation for the pairs of short mesodermal clones accompanied by long or short neural clones observed in the mouse retrospective analyses (Tzouanacou et al., 2009). Long clones in both tissues can be generated by the other source of progenitors of dual fate: the NMPZ located at the interface between the specialised axial stem zones. These are contained in an arc 5-10 cell diameters thin, containing about 200 cells (see Figure 5a-c and Supplementary Videos S8 and S11). Most of these cells remain in this domain during the 15 hours of elongation examined here, and a few of these produce daughters in different domains. However, all cells in the NMPZ domain are evenly mingled, irrespective of their fate; they are also equally multipotent and can become part of either the mesodermal or the neural stem zone (see Figure 6).

What mechanisms drive cells in the NMPZ to adopt different fates? We found no evidence for over-proliferation or oriented divisions in the NMPZ (see Figure 5 and Video S13). Another hypothesis is that daughters of cells in the NMPZ differ in their response to signals from adjacent territories, and move actively into either the mesodermal or neural stem zone. This might be due to variations in the expression of pro-mesodermal and pro-neural genes (see Introduction); this hypothesis could not be tested, and also presupposes that the cells could exert directional movements within the epiblast, a tight epithelium. We propose a simpler mechanism, which does not require any of these assumptions and only relies on the general properties of the tight epithelium the MSZ, NMPZ and NSZ are part of. The divergent collective flows the cells surrounding the NMPZ (see Figure 6b and Video S14) involve thousands of cells in the mesodermal and neural fields, compared with a few hundred in the intervening NMPZ; they also increase in magnitude away from the NMPZ. They cannot therefore be caused by the NMPZ. Instead, the retraction of the node (the shortening of the PS) and the convergence and extension of the neural plate are likely to produce shear between the MSZ and NSZ. The NMPZ cells disperse locally and isotropically after cell division, and we propose that they are then pulled passively by the massive, collective movements surrounding their domain. The descendants of the NMPZ cells only assume a stable identity once they reach the neural or mesodermal domain.

What role do the NMP cells play in vivo? Our results show that most cells elongating the axis are born directly in the specialized stem zones (mesodermal and neural), and that the neuromesodermal zone contributes very few cells to either of these; this might explain why their ablation in mouse only produces limited defects (Mugele et al., 2018). It also appears that bipotential progenitors do not actively allocate cells to the specialized axial stem zones based on intrinsic determinants, and do not control the balance of mesoderm and neural tissue produced. Therefore, the neuromesodermal state does not seem to be an obligatory step in populating most of axial stem zones or in fuelling axial elongation. However, the NMPZ functioning reveals a principle of how two committed domains (stem zones) can be separated in the absence of a classical boundary. The results suggest that, in embryos of regulative development, a region of bipotential cells is created at their interface. This prevents each of the committed domains from drawing cells from the other, instead pulling from the reservoir of cells that can adopt their identity.

## Methods

Fertilized eggs of wild-type hens (Bovans Brown) were obtained from WinterEgg Farm (Thriplow, Herts, UK), and of transgenic myr-EGFP birds (Rozbicki et al., 2015) from the Roslin Institute (Royal (Dick) School of Veterinary Studies, University of Edinburgh, Roslin, UK).

### Embryo culture

Fresh, fertilised hen’s eggs were kept for up to one week at 15°C until use, and incubated at 38°C for embryo to reach the desired stages. Embryos were explanted and manipulated in Pannett-Compton saline (Pannett and Compton, 1924), and cultured using New’s technique (New, 1955), as modified and described previously (Voiculescu et al., 2008).

### Fluorescent labelling

Working dilutions of lipophilic CM-DiI and DiO (Invitrogen, cat. # C7001 and D1125) were prepared fresh, by adding 1 μl of stock solutions (0.5% w/v in ethanol, kept at −20°C) to 9 μl of aqueous sucrose solution (6% w/v) pre-warmed to 37°C. Finely drawn capillaries, with a tip broken to a bore of about 1-2 μm, were used to deposit warm solution close to the epiblast of embryos kept in saline.

### Whole node transplantations

Donor embryos were fluorescently labelled by CMFDA (Invitrogen, cat. # C2925) by bathing whole, for 1 hour at 38°C, in Pannett-Compton solution containing CMFDA diluted 1:250 from stock solution (10 mM in DMSO, kept at −20°C). They were then transferred three times for 10 minutes each in fresh Pannett-Compton saline to remove excess dye, before having their nodes excised. The nodes were excised from unlabelled receiving embryos, prepared in New culture; the labelled nodes were aspirated and transferred with the aid of a P2 pipette adjusted to 0.2 μl.

### Electroporations

Electroporation of epiblast of donor embryos (Figure 6c) was done with the 3’-fluorescein tagged standard control morpholino (Gene-Tools), diluted 1:5 from the aqueous stock (1 mM) in a solution containing 6% (w/v) sucrose and plasmid DNA (100 μg ml-1) (Voiculescu and Stern, 2017; Voiculescu et al., 2008). A train of three square pulses of 5.7 V, 50 ms duration at 500 ms interval was applied using an Intracel TSS 20 Ovodyne electroporator.

### Microsurgery

Surgical manipulations were done using fine tungsten needles sharpened by electroelution (Brady, 1965). For RNA-seq (Figure 1h), the region containing the resident, fluorescently-labelled cells in the mature organizer was first excised whole from the embryo, which was then subdivided into the brightly fluorescent regions of the anterior primitive streak and the faintly- or non-labelled regions. For the transplantation experiments exemplified in Figure 6c-g, incisions were made in the epiblast of the electroporated donors, following the contours of the region shown in Figure 6c. The cut epiblast was then gently separated from the embryo by scraping under it to separate the underlying mesoderm. The excised material was then cut into four pieces, each of which transplanted into the regions indicated in Figure 6d-f. In the receiving embryos, mounted in New’s culture rings (endoderm-side up), a keyhole incision was made over the intended grafting area, and a small cut was made in the epiblast. The grafted material was pushed in, and covered with the lower layers of the receiving embryo to help healing. Operated embryos were kept at room temperature for 2-4 hours, conditions under which embryos are in developmental diapause but healing can take place, and then placed in a humidified box at 38°C and allowed to grow.

### Gene expression analyses

RT-PCR (Figure 1j) was done using Phusion High-Fidelity (ThermoFisher, cat. # F553L) a cDNA generated from 5 nodes dissected at stages HH 4-7. The list of primers used The PCR fragments were gel-purified and sequenced to check their sequence corresponds to the intended transcript, and DIG-RNA probes were synthesised using the SP6 promoter included in the reverse primers, as described in GEISHA. In situ hybridizations were performed using the methods described earlier (Streit and Stern, 2001)․.

### RNA-seq

Total RNA was column-extracted from freshly dissected or laser-captured (LCM) material using RNeasy Micro Kit (Qiagen, cat. # 74004), with a yield of 1-8 ng per sample. Several fixation methods were compared, of which we found Methacarn (methanol:chloroform:acetic acid, 6:3:1 volume proportions) to be most suitable in terms of morphology preservation and quality of extracted RNA. Embryos with the mature organizer fluorescently labelled were grown in vitro (see above). At the end of incubation, the glass ring with vitelline membrane supporting the embryo was lifted from the culture dish, the membrane and embryo quickly washed with Pannett-Compton solution, and placed in a watch glass over a small pool of ice-cold fixative; fixative was also used to flood the inside of the glass ring and submerge the embryo. The assembly was kept on ice for 30 minutes. Dissected embryos were brought into absolute ethanol, embedded in paraffin and serially cut at 5 μm. Sections were placed on metal framed, 0.9 μm thick POL-membranes (Leica, REF 11505191), and the regions of interest were dissected using a Leica LMD 600 laser capture microdissection system fitted with a x63 lens and fluorescence optics. The total RNA was quality-checked on a pico chip (Bioanalyzer 2100, Agilent), amplified with SMARTer kit (Clontech), and sequenced on an Illumina HiSeq 4000 platform (100bp runs, pair end).

Gene counts and comparisons of gene expression levels between each pair of samples were performed using the Trapnell pipeline (Trapnell et al., 2012). For each pair, we assigned a score of 1 to genes at least 5 times more expressed in the “red” sample (from the MSZ) than in the “white” one (from surrounding regions), and a score of 2 if there was a significant gene count in the “red” sample but zero in the “white” one. Genes were then ranked based on the aggregate score (sheet 1 in Supplementary Table S13, sheet 1). For further assessment by RT-PCR (Supplementary Table S13, sheet 2), we discarded the genes whose expression patterns were known not to be specific to the regions of interest, but included low-ranking genes showing very dissimilar levels in at least some pairs of samples.

### Time-lapse imaging and processing

#### Microscopy

Time-lapse epifluorescence movies were acquired with a PlanApo N 2x/0.08 lens on an Olympus IX71 wide-field microscope, fitted with fluorescence excitation source (CoolLed pE-2) and appropriate emission optics, motorised stage (Prior), and Hamamatsu C8484 camera controlled by HCImage software. The system was enclosed in a Perspex box, one wall of which is the top of a Marsh Automatic Incubator (Lyon, U.S.A.) to provide thermoregulation and air circulation.

For multi-fluorescence imaging, transgenic embryos were mounted in custom imaging dishes (Voiculescu and Stern, 2012), and placed under an upright 2-photon microscope (LaVision BioTech TriM Scope II) with the fs laser operating at 915 nm. The 2-photon microscope was equipped with thermo-regulated hood, and imaging performed using a 25x/1.01 water dipping lens (Olympus XLPLN25XWMP2). Immersol W 2010 (Zeiss, cat. # 444969-0000-000) was used as a medium between the objective and the imaging dish. Two partially overlapping fields (400 μm x 400 μm) were acquired to cover the anterior primitive streak and surrounding regions; acquisition fields were reassigned as needed, to keep the regressing node in the centre of one field. For each field, 90-100 optical planes (1282 x 1282 xy resolution, 16-bit depth) were acquired with a 1.5 μm z-spacing to encompass the entire thickness of the embryo.

#### Preparation of multiphoton raw imaging data

The two fields acquired per time point were stitched using the Fiji Stitching Plugin, and then registered using the Multiview Reconstruction plugin in Fiji, working in the hdf5 container image format. Initially the time points were manually translated for rough alignment, followed by an automated rigid body registration with normalization. This was then output without any cropping of the data into tiff image stacks for use in cell tracking within FIJI and RACE.

#### Manual tracking

Tracking and production of lineage trees were accomplished using the FIJI package with the TrackMate plugin (Tinevez et al., 2017), and this was also used to analyse the distribution of lineages with different numbers of cell divisions (Figure 5e, f). Manual tracks were independently drawn and checked by two members of the team. A bespoke XML parser was written to import the tracking file into MATLAB for further analyses.

#### Automatic Segmentation

An automatic segmentation pipeline was developed on the basis of the RACE algorithm (Stegmaier et al., 2016) and implemented as a custom pipeline in the open-source image analysis software XPIWIT (Bartschat et al., 2016). Initially, image noise was reduced using a 3D median filter with a radius of 1 voxel (3×3×3 filter mask) followed by a Hessian-based objectness filter that emphasized plane-like structures in the 3D images (for details, see (Antiga, 2007; Bartschat et al., 2016)). The enhanced cell walls were then binarized using a manually adjusted threshold and converted to a distance map where intensity values of the image represent the Euclidean distance in voxels to the closest foreground voxel of the binarized cell walls (bright spots in the centres of the cells, darker towards the cell membranes). The distance map was then multiplied with the inverted raw image, to further lower the intensity of the distance map on the cell walls. Finally, we applied a morphological watershed algorithm (Beare and Lehmann, 2006) in 3D on the enhanced and inverted Euclidean distance map image (dark valleys in cell interior and bright boundaries on the cell walls) to obtain the final segmentation. This procedure was repeated separately for each of the unregistered 3D raw images and the segmented label images were subsequently used for tracking.

#### Registration and Tracking

To compensate for the posterior retraction movements of the stem zones regions occurring during acquisition, we initially preregistered the 3D images semi-automatically using a custom-made MATLAB user interface. Next, we performed an elastic registration using the B-spline method implemented in elastix (Klein et al., 2010) to obtain a displacement map for the entire 3D image that was used for improved tracking. Tracking was implemented in MATLAB and we identified corresponding objects in a backwards fashion (i.e., from the last to the first time point) by identifying the maximally overlapping connected components in adjacent frames. For each of the segments present in t+1, we identified the segment with the largest overlap in time point t and remembered this predecessor id for the subsequent reconstruction of the object trajectories. If the maximally overlapping segment was the background (e.g., if the cell moved out of the field of view), the traceback was stopped. To improve the overlap between consecutive frames, we kept one of the frames as a reference (t+1) and then transformed the respective label image of the neighbouring frame t using the displacement map identified during the elastic registration of the raw images with a nearest neighbour interpolation scheme to avoid distorting the discrete segment labels. Note that the elastic registration was only used for the identification of the maximally overlapping segments in neighbouring frames and that the final positions used for all visualizations correspond to the locations of the shift-corrected images.

#### Visualization and data analyses

Visualization and data analyses were performed in MATLAB. Raw image slices were extracted from the 3D stacks by fitting a 2D surface to the locations of manually tracked objects at each of the time points. We then extracted intensity values from the original stacks based on the estimated surface height at each of the voxel locations and performed a contrast-limited adaptive histogram equalization (Zuiderveld, 1994) for improved visibility and contrast of the extracted image slice. For improved visualization and interpretability, all manual and automatic trajectories were smoothed using a moving average filter with a span of 11 time points. Moreover, the node position was manually selected in all of the frames and was used as a point of reference, i.e., all images and cell locations were translated with respect to this point, such that the node remained static throughout the sequence and the surrounding tissue moved relative to this point.

## Acknowledgements

This work was funded by the Wellcome Trust (RCDF 088380/09/Z to O.V.), the German Research Foundation (DFG, Grant No. MI 1315/4 associated with SPP 1736 Algorithms for Big Data to J.S.) and the Helmholtz Association (BioInterfaces in Technology and Medicine Program, R.M.). A.K. was supported by the Carlsberg Foundation, and I.K. by a Wellcome Trust doctoral fellowship (097416/Z/11/Z). We thank the High-Throughput Genomics Group at the Wellcome Trust Centre for Human Genetics (funded by Wellcome Trust grant reference 090532/Z/09/Z and MRC Hub grant G0900747 91070) for the generation of the Sequencing data, and Kevin O’Holleran (Cambridge Advanced Imaging Centre) for assistance with the multi-photon microscope. We are grateful to Alfonso Martinez-Arias, Claudio D. Stern, Ben Steventon and Anestis Tsakiridis for comments on the early draft of the manuscript.

## Author contributions

T.R.W., I.K. and O.V. performed time-lapse microscopy and microdissection, constructed RNA-seq libraries and analysed the deep-sequencing data. A.K. and O.V. did the manual tracking and annotation, and J.S. the automatic segmentation, tracking and wrote the MATLAB scripts. O.V. wrote the manuscript, with input from all authors.

## Materials & Correspondence

Requests should be addressed to Octavian Voiculescu

## Supplementary information

**Video S1. The organizer becomes a stem zone at the end of gastrulation.**

The organizer and the middle of the primitive streak fluorescently labelled with CM-DiI (red) at the end of gastrulation (HH 3^+^). Time is indicated in hours:minutes.

**Video S2. Cells around the mature organizer do not ingress, instead contributing to the neural plate.**

The tip, middle and posterior of the primitive streak labelled with DiO (green), and the epiblast left to these locations labelled with CM-DiI (red).

**Table S3. Gene counts in the mature organizer (R) and surrounding tissues (W).**

L and M correspond to the method of isolation employed (laser capture or manual dissection).

**Table S4. Primers used to for RT-PCR and in situ hybridisation probes.**

Genes examined by in situ hybridization are highlighted in yellow.

**Video S5. Multi-photon imaging of the regions involved in axial elongation, in membrane-EGFP transgenic embryos.**

The scanned regions include and surround the mature organizer and the portions of the streak; acquisition fields were changed to follow the node during its posterior retraction.

**Video S6. Stereo-pair movie of the regions fuelling axial elongation.**

Frames are registered with respect to the node to compensate for its posterior retraction.

**Video S7. Comparison between the manually and automatically tracked cells.**

Lineages traced manually (green) are superimposed on the trajectories of automatically segmented and tracked cells (red). Only a fifth of the latter are displayed, for clarity.

**Video S8. Stereo movie of axial stem cell lineages, coloured by fate.**

Cells of neural fate in blue, those contributing to the anterior quadrant and lateral sides of the mature organizer in yellow and red, respectively. Magenta denotes cells of the zone of mixed, neural and mesodermal precursors (ZNMP). Cells pulled towards, and later ingressing through the primitive streak, are coloured green and turquoise.

**Video S9. Cell movements of the axial stem cells**

The actual position of each cell is indicated by a dot, coloured as in Video S8. As cells move, their trajectories over the previous 3 hours are displayed as fading tails. Left, lineages displayed against the raw images’ projection to provide anatomical context. Right, only the tracked lineages are displayed.

**Video S10. Ingression events occur outside the primitive streak, including the neural plate.**

Neural plate cells are shown in blue, ZNMP cells in magenta, and cells contributing to any level of the primitive streak in green. Ingression events are marked by an enlarged dot, at the time when they occur.

**Video S11. Cell divisions in the ZNMP producing descendants of different fate.**

White bars connect daughters of cell divisions of cells in the ZNMP (magenta) that adopt different fates. Other cell types are coloured as in Movies S8, S9 and S10.

**Video S12. Regional distribution of lineages with different cycling times.**

Lineages are colour-coded with respect to the number of cell divisions observed during the duration of the movie (see lower middle inset for key). Nine-hour trajectories are displayed.

**Video S13. Angles of cell division and dispersion of daughter cells.**

Cells are colour-coded with respect to their fate: neural plate (blue), ZNMP (magenta) and primitive streak-destined (green), as in as in Movies S8, S9, S10 and S11. Left, white bars connect daughter cells after each cell division. Middle column, angle of cell divisions with respect to mother cell trajectory. Right column, distance between sister cells following cell division.

**Video S14. Collective cell flows around the zone of multiple precursors.**

Left, ZNMP cells coloured in magenta; cells destined for the neural plate (blue), mesodermal stem zone (yellow and red) and the more caudal primitive streak (green) are shown with fading 6 hour-long trails. Trails of automatically segmented cells are also shown in grey levels, to illustrate the cell flows with higher density.

Right, the cells of different fates are shown with the same colour scheme without trails, against the imaged regions to provide anatomical context at each time point.

## References

Acloque, H., Ocana, O.H., Matheu, A., Rizzoti, K., Wise, C., Lovell-Badge, R., and Nieto, M.A. (2011). Reciprocal repression between Sox3 and snail transcription factors defines embryonic territories at gastrulation. Dev Cell 21, 546–558.

Acloque, H., Ocana, O.H., and Nieto, M.A. (2012). Mutual exclusion of transcription factors and cell behaviour in the definition of vertebrate embryonic territories. Curr Opin Genet Dev 22, 308–314.

Acloque, H., Ocana, O.H., Abad, D., Stern, C.D., and Nieto, M.A. (2017). Snail2 and Zeb2 repress P-cadherin to define embryonic territories in the chick embryo. Development 144, 649–656.

Aires, R., Dias, A., and Mallo, M. (2018). Deconstructing the molecular mechanisms shaping the vertebrate body plan. Curr Opin Cell Biol 55, 81–86.

Anderson, C., Khan, M.A., Wong, F., Solovieva, T., Oliveira, N.M., Baldock, R.A., Tickle, C., Burt, D.W., and Stern, C.D. (2016). A strategy to discover new organizers identifies a putative heart organizer. Nat Commun 7, 12656.

Antiga, L. (2007). Generalizing vesselness with respect to dimensionality and shape. Insight J. 3, 1–14.

Attardi, A., Fulton, T., Florescu, M., Shah, G., Muresan, L., Lenz, M.O., Lancaster, C., Huisken, J., Oudenaarden, A. van, and Steventon, B. (2018). Neuromesodermal progenitors are a conserved source of spinal cord with divergent growth dynamics. Development 145, dev166728.

Bartschat, A., Hubner, E., Reischl, M., Mikut, R., and Stegmaier, J. (2016). XPIWIT--an XML pipeline wrapper for the Insight Toolkit. Bioinformatics 32, 315–317.

Beare, R., and Lehmann, G. (2006). The watershed transform in ITK-discussion and new developments. Insight J 92, 1–24.

Brady, J. (1965). A simple technique for making very fine, durable dissecting needles by sharpening tungsten wire electrolytically. Bull. World Health Organ. 32, 143–144.

Brown, J.M., and Storey, K.G. (2000). A region of the vertebrate neural plate in which neighbouring cells can adopt neural or epidermal fates. Curr Biol 10, 869–872.

Cambray, N., and Wilson, V. (2002). Axial progenitors with extensive potency are localised to the mouse chordoneural hinge. Development 129, 4855–4866.

Cambray, N., and Wilson, V. (2007). Two distinct sources for a population of maturing axial progenitors. Development 134, 2829–2840.

Catala, M., Teillet, M.A., and Le Douarin, N.M. (1995). Organization and development of the tail bud analyzed with the quail-chick chimaera system. Mech Dev 51, 51–65.

Davis, R.L., and Kirschner, M.W. (2000). The fate of cells in the tailbud of Xenopus laevis. Development 127, 255–267.

Denham, M., Hasegawa, K., Menheniott, T., Rollo, B., Zhang, D., Hough, S., Alshawaf, A., Febbraro, F., Ighaniyan, S., Leung, J., et al. (2015). Multipotent caudal neural progenitors derived from human pluripotent stem cells that give rise to lineages of the central and peripheral nervous system. Stem Cells 33, 1759–1770.

Frith, T.J., Granata, I., Wind, M., Stout, E., Thompson, O., Neumann, K., Stavish, D., Heath, P.R., Ortmann, D., Hackland, J.O., et al. (2018). Human axial progenitors generate trunk neural crest cells in vitro. Elife 7.

Garriock, R.J., Chalamalasetty, R.B., Kennedy, M.W., Canizales, L.C., Lewandoski, M., and Yamaguchi, T.P. (2015). Lineage tracing of neuromesodermal progenitors reveals novel Wnt-dependent roles in trunk progenitor cell maintenance and differentiation. Development 142, 1628–1638.

Gont, L.K., Steinbeisser, H., Blumberg, B., and de Robertis, E.M. (1993). Tail formation as a continuation of gastrulation: the multiple cell populations of the Xenopus tailbud derive from the late blastopore lip. Development 119, 991–1004.

Gouti, M., Tsakiridis, A., Wymeersch, F.J., Huang, Y., Kleinjung, J., Wilson, V., and Briscoe, J. (2014). In vitro generation of neuromesodermal progenitors reveals distinct roles for wnt signalling in the specification of spinal cord and paraxial mesoderm identity. PLoS Biol 12, e1001937.

Gouti, M., Delile, J., Stamataki, D., Wymeersch, F.J., Huang, Y., Kleinjung, J., Wilson, V., and Briscoe, J. (2017). A Gene Regulatory Network Balances Neural and Mesoderm Specification during Vertebrate Trunk Development. Dev Cell 41, 243–261 e7.

de Graaf, C.A., and Metcalf, D. (2011). Thrombopoietin and hematopoietic stem cells. Cell Cycle 10, 1582–1589.

Gros, J., Feistel, K., Viebahn, C., Blum, M., and Tabin, C.J. (2009). Cell movements at Hensen’s node establish left/right asymmetric gene expression in the chick. Science 324, 941–944.

Henrique, D., Abranches, E., Verrier, L., and Storey, K.G. (2015). Neuromesodermal progenitors and the making of the spinal cord. Development 142, 2864–2875.

Hitchcock, I.S., and Kaushansky, K. (2014). Thrombopoietin from beginning to end. Br J Haematol 165, 259–268.

Holmdahl, D.E. (1925). Experimentelle Untersuchungen über die Lage der Grenze zwischen primärer und secondärer Körperentwicklung beim Huhn. Anat. Anz. 59, 393–396.

Holmdahl, D.E. (1939). Die Morphogenese des Vertebratorganismus vom formalen und experimentellen Gesichtspunkt. Wilhelm Roux Arch Entwickl Mech Org 139, 191–226.

Hsieh, J.C., Kodjabachian, L., Rebbert, M.L., Rattner, A., Smallwood, P.M., Samos, C.H., Nusse, R., Dawid, I.B., and Nathans, J. (1999). A new secreted protein that binds to Wnt proteins and inhibits their activities. Nature 398, 431–436.

Iimura, T., Yang, X., Weijer, C.J., and Pourquie, O. (2007). Dual mode of paraxial mesoderm formation during chick gastrulation. Proc Natl Acad Sci U A 104, 2744–2749.

Javali, A., Misra, A., Leonavicius, K., Acharya, D., Vyas, B., and Sambasivan, R. (2017). Co-expression of Tbx6 and Sox2 identifies a novel transient neuromesoderm progenitor cell state. Development.

Joubin, K., and Stern, C.D. (1999). Molecular interactions continuously define the organizer during the cell movements of gastrulation. Cell 98, 559–571.

Kashuba, E., Kashuba, V., Sandalova, T., Klein, G., and Szekely, L. (2002). Epstein-Barr virus encoded nuclear protein EBNA-3 binds a novel human uridine kinase/uracil phosphoribosyltransferase. BMC Cell Biol 3, 23.

Kaushansky, K. (2016). Thrombopoietin and its receptor in normal and neoplastic hematopoiesis. Thromb J 14, 40.

Kimelman, D. (2016). Tales of Tails (and Trunks): Forming the Posterior Body in Vertebrate Embryos. Curr Top Dev Biol 116, 517–536.

Kinder, S.J., Tsang, T.E., Wakamiya, M., Sasaki, H., Behringer, R.R., Nagy, A., and Tam, P.P.L. (2001). The organizer of the mouse gastrula is composed of a dynamic population of progenitor cells for the axial mesoderm. Development 128, 3623–3634.

Klein, S., Staring, M., Murphy, K., Viergever, M.A., and Pluim, J.P. (2010). Elastix: a toolbox for intensity-based medical image registration. IEEE Trans. Med. Imaging 29, 196–205.

Koch, F., Scholze, M., Wittler, L., Schifferl, D., Sudheer, S., Grote, P., Timmermann, B., Macura, K., and Herrmann, B.G. (2017). Antagonistic Activities of Sox2 and Brachyury Control the Fate Choice of Neuro-Mesodermal Progenitors. Dev Cell 42, 514–526 e7.

Le Douarin, N.M., Teillet, M.A., and Catala, M. (1998). Neurulation in amniote vertebrates: a novel view deduced from the use of quail-chick chimeras. Int J Dev Biol 42, 909–916.

Lippmann, E.S., Williams, C.E., Ruhl, D.A., Estevez-Silva, M.C., Chapman, E.R., Coon, J.J., and Ashton, R.S. (2015). Deterministic HOX patterning in human pluripotent stem cell-derived neuroectoderm. Stem Cell Rep. 4, 632–644.

Martin, B.L., and Kimelman, D. (2009). Wnt signaling and the evolution of embryonic posterior development. Curr Biol 19, R215–9.

Martin, B.L., and Kimelman, D. (2012). Canonical Wnt signaling dynamically controls multiple stem cell fate decisions during vertebrate body formation. Dev Cell 22, 223–232.

Mathis, L., Kulesa, P.M., and Fraser, S.E. (2001). FGF receptor signalling is required to maintain neural progenitors during Hensen’s node progression. Nat Cell Biol 3, 559–566.

McDole, K., Guignard, L., Amat, F., Berger, A., Malandain, G., Royer, L.A., Turaga, S.C., Branson, K., and Keller, P.J. (2018). In Toto Imaging and Reconstruction of Post-Implantation Mouse Development at the Single-Cell Level. Cell 175, 859–876.e33.

Mendes, R.V., Martins, G.G., Cristovao, A.M., and Saude, L. (2014). N-cadherin locks left-right asymmetry by ending the leftward movement of Hensen’s node cells. Dev Cell 30, 353–360.

Michaille, J.J., Tili, E., Calin, G.A., Garin, J., Louwagie, M., and Croce, C.M. (2005). Cloning and characterization of cDNAs expressed during chick development and encoding different isoforms of a putative zinc finger transcriptional regulator. Biochimie 87, 939–949.

Mugele, D., Moulding, D., Savery, D., Mole, M., Greene, N., Martinez-Barbera, J.P., and Copp, A. (2018). Genetic approaches in mice demonstrate that neuro-mesodermal progenitors express T/Brachyury but not Sox2. BioRxiv 503854.

New, D.A.T. (1955). A New Technique for the Cultivation of the Chick Embryo in vitro. J. Embryol. Exp. Morphol. 3, 326–331.

Pannett, C., and Compton, A. (1924). The Cultivation of Tissues in Saline Embryonic Juice. The Lancet 203, 381–384.

Patten, I., Kulesa, P., Shen, M.M., Fraser, S., and Placzek, M. (2003). Distinct modes of floor plate induction in the chick embryo. Development 130, 4809–4821.

Psychoyos, D., and Stern, C.D. (1996). Fates and migratory routes of primitive streak cells in the chick embryo. Development 122, 1523–1534.

Rozbicki, E., Chuai, M., Karjalainen, A.I., Song, F., Sang, H.M., Martin, R., Knolker, H.J., MacDonald, M.P., and Weijer, C.J. (2015). Myosin-II-mediated cell shape changes and cell intercalation contribute to primitive streak formation. Nat Cell Biol 17, 397–408.

Selleck, M.A., and Stern, C.D. (1991). Fate mapping and cell lineage analysis of Hensen’s node in the chick embryo. Development 112, 615–626.

Selleck, M.A.J., and Stern, C.D. (1992a). Evidence for Stem Cells in the Mesoderm of Hensen’s Node and Their Role in Embryonic Pattern Formation (Springer, Boston, MA: NATO ASI Series (Series A: Life Sciences)).

Selleck, M.A.J., and Stern, C.D. (1992b). COMMITMENT OF MESODERM CELLS IN HENSENS NODE OF THE CHICK-EMBRYO TO NOTOCHORD AND SOMITE. Development 114, 403–415.

Sheng, G., dos Reis, M., and Stern, C.D. (2003). Churchill, a zinc finger transcriptional activator, regulates the transition between gastrulation and neurulation. Cell 115, 603–613.

Stegmaier, J., Amat, F., Lemon, W.C., McDole, K., Wan, Y., Teodoro, G., Mikut, R., and Keller, P.J. (2016). Real-Time Three-Dimensional Cell Segmentation in Large-Scale Microscopy Data of Developing Embryos. Dev Cell 36, 225–240.

Streit, A., and Stern, C.D. (2001). Combined whole-mount in situ hybridization and immunohistochemistry in avian embryos. Methods 23, 339–344.

Takemoto, T., Uchikawa, M., Yoshida, M., Bell, D.M., Lovell-Badge, R., Papaioannou, V.E., and Kondoh, H. (2011). Tbx6-dependent Sox2 regulation determines neural or mesodermal fate in axial stem cells. Nature 470, 394–398.

Tanaka, T., Urade, Y., Kimura, H., Eguchi, N., Nishikawa, A., and Hayaishi, O. (1997). Lipocalin-type prostaglandin D synthase (beta-trace) is a newly recognized type of retinoid transporter. J Biol Chem 272, 15789–15795.

Taniguchi, Y., Kurth, T., Weiche, S., Reichelt, S., Tazaki, A., Perike, S., Kappert, V., and Epperlein, H.H. (2017). The posterior neural plate in axolotl gives rise to neural tube or turns anteriorly to form somites of the tail and posterior trunk. Dev Biol 422, 155–170.

Tinevez, J.Y., Perry, N., Schindelin, J., Hoopes, G.M., Reynolds, G.D., Laplantine, E., Bednarek, S.Y., Shorte, S.L., and Eliceiri, K.W. (2017). TrackMate: An open and extensible platform for single-particle tracking. Methods 115, 80–90.

Trapnell, C., Roberts, A., Goff, L., Pertea, G., Kim, D., Kelley, D.R., Pimentel, H., Salzberg, S.L., Rinn, J.L., and Pachter, L. (2012). Differential gene and transcript expression analysis of RNA-seq experiments with TopHat and Cufflinks. Nat Protoc 7, 562–578.

Tsakiridis, A., Huang, Y., Blin, G., Skylaki, S., Wymeersch, F., Osorno, R., Economou, C., Karagianni, E., Zhao, S., Lowell, S., et al. (2014). Distinct Wnt-driven primitive streak-like populations reflect in vivo lineage precursors. Development 141, 1209–1221.

Turner, D.A., Rue, P., Mackenzie, J.P., Davies, E., and Martinez Arias, A. (2014). Brachyury cooperates with Wnt/beta-catenin signalling to elicit primitive-streak-like behaviour in differentiating mouse embryonic stem cells. BMC Biol 12, 63.

Tzouanacou, E., Wegener, A., Wymeersch, F.J., Wilson, V., and Nicolas, J.F. (2009). Redefining the progression of lineage segregations during mammalian embryogenesis by clonal analysis. Dev Cell 17, 365–376.

Verrier, L., Davidson, L., Gierliński, M., Dady, A., and Storey, K.G. (2018). Neural differentiation, selection and transcriptomic profiling of human neuromesodermal progenitor-like cells in vitro. Development 145, dev166215.

Vogt, H. (1926). Über Wachstum und Gestaltungbewegungen am hinteren Körperende der Amphibien. Anat. Anz. 61, 62–75.

Voiculescu, O., and Stern, C.D. (2012). Assembly of imaging chambers and high-resolution imaging of early chick embryos. Cold Spring Harb Protoc 2012.

Voiculescu, O., and Stern, C.D. (2017). Manipulating Gene Expression in the Chick Embryo. Methods Mol Biol 1565, 105–114.

Voiculescu, O., Papanayotou, C., and Stern, C.D. (2008). Spatially and temporally controlled electroporation of early chick embryos. Nat Protoc 3, 419–426.

Voiculescu, O., Bodenstein, L., Lau, I.J., and Stern, C.D. (2014). Local cell interactions and self-amplifying individual cell ingression drive amniote gastrulation. Elife 3, e01817.

Wilson, V., Olivera-Martinez, I., and Storey, K.G. (2009). Stem cells, signals and vertebrate body axis extension. Development 136, 1591–1604.

Wymeersch, F.J., Huang, Y., Blin, G., Cambray, N., Wilkie, R., Wong, F.C., and Wilson, V. (2016). Position-dependent plasticity of distinct progenitor types in the primitive streak. Elife 5, e10042.

Zhang, A., Dong, Z., and Yang, T. (2006). Prostaglandin D2 inhibits TGF-beta1-induced epithelial-to-mesenchymal transition in MDCK cells. Am J Physiol Ren. Physiol 291, F1332–42.

Zuiderveld, K. (1994). Contrast limited adaptive histogram equalization. (Academic Press Professional Inc.), pp. 474–485.

